# Time perception reflects individual differences in motor and non-motor symptoms of Parkinson’s disease

**DOI:** 10.1101/2023.03.02.530411

**Authors:** Emily DiMarco, Renata Sadibolova, Angela Jiang, Brittany Liebenow, Rachel E. Jones, Ihtsham ul Haq, Mustafa S. Siddiqui, Devin B. Terhune, Kenneth T. Kishida

## Abstract

Dopaminergic signaling in the striatum has been shown to play a critical role in the perception of time. Decreasing striatal dopamine efficacy is at the core of Parkinson’s disease (PD) motor symptoms and changes in dopaminergic action have been associated with many comorbid non-motor symptoms in PD. We hypothesize that patients with PD perceive time differently and in accordance with their specific comorbid non-motor symptoms and clinical state. We recruited patients with PD and compared individual differences in patients’ clinical features with their ability to judge millisecond to second intervals of time (500ms-1100ms) while on or off their prescribed dopaminergic medications. We show that individual differences in comorbid non-motor symptoms, PD duration, and prescribed dopaminergic pharmacotherapeutics account for individual differences in time perception performance. We report that comorbid impulse control disorder is associated with temporal overestimation; depression is associated with decreased temporal accuracy; and PD disease duration and prescribed levodopa monotherapy are associated with reduced temporal precision and accuracy. Observed differences in time perception are consistent with hypothesized dopaminergic mechanisms thought to underlie the respective motor and non-motor symptoms in PD, but also raise questions about specific dopaminergic mechanisms. In future work, time perception tasks like the one used here, may provide translational or reverse translational utility in investigations aimed at disentangling neural and cognitive systems underlying PD symptom etiology.

**One Sentence Summary:** Quantitative characterization of time perception behavior reflects individual differences in Parkinson’s disease motor and non-motor symptom clinical presentation that are consistent with hypothesized neural and cognitive mechanisms.

## INTRODUCTION

Time perception is a fundamental cognitive process that intersects with basic human cognitive functions, such as attention, memory, sensorimotor processing, and decision-making (1–6). Time perception ranging from millisecond to minute durations is referred to as interval timing, and studies utilizing pharmacologic, genetic, neuroimaging, and stimulation-based manipulations widely support the involvement of striatal dopamine in this process (7–18). Notably, investigations into dopaminergic disorders in patient populations (e.g. Schizophrenia and Parkinson’s disease) show specific quantifiable changes in interval timing behavior. For example, Schizophrenia (hypothesized to reflect a hyperdopaminergic state) is associated with overestimation of time intervals (19–21) and Parkinson’s disease (a hypodopaminergic state) has been associated with reduced ability to discriminate between time intervals (22,23).

Parkinson’s disease (PD) is caused by the irreversible loss of midbrain dopamine neuron terminals, which is associated with significant motor deficits (24,25). However, patients with PD also experience significant non-motor symptoms in the form of neuropsychiatric, autonomic, sleep, and sensory changes (26). Many of these comorbid conditions are hypothesized to have some form of dopaminergic etiology, but the precise mechanisms are not well understood. For example, Impulse Control Disorder (ICD) is a behavioral addiction characterized by the need to perform pleasurable and often risky behaviors compulsively and repetitively, which can be caused and/or intensified by certain dopaminergic therapies (DT) prescribed to alleviate PD motor symptoms (27–31). It is hypothesized that patients with ICD experience a state akin to a hyperdopaminergic state based on the link between ICD induction by dopamine receptor agonists with a preferential affinity for dopamine (D3) receptors in the ventral striatum (31,32). Additionally, depression affects approximately 40% of patients with PD (33,34) – nearly twice the rate of the general population. Depression in PD increases with PD symptom severity, physical disability, and PD duration (34). The etiology of depression remains unclear; however, substantial evidence implicates dopamine dysfunction in affective disorders (35,36). In particular, diminished Ventral Tegmental Area (VTA) dopaminergic activity and reduced VTA-striatal connectivity have been linked to anhedonia and amotivation (36,37), common depressive symptoms seen in patients with PD.

Generally, time perception in patient cohorts appears to be altered in a manner consistent with the hypothesized roles of dopamine in ICD (31,32) and depression (35–37). Impulsive patients in populations outside of patients with PD, such as schizophrenia and borderline personality disorder, tend to present with increased accuracy and precision variability on interval timing tasks (38,39). Patients with depressive symptoms have been observed to underestimate the duration of time intervals (40–42). Despite the increased rate of comorbidity in PD, the impact of comorbid depression or ICD on interval timing and the underlying neurobiology leading to depression and ICD in PD remains poorly understood.

Additional individual differences in the clinical state of patients with PD may be inferred from the strategies most effective at managing individual patients’ symptoms with prescribed medications. The medications used to treat PD motor symptoms primarily target the dopaminergic system (43,44). These DT affect the dopaminergic system in different ways, are prescribed according to patients’ specific symptom management needs, and are often required to be increased or changed in response to disease progression or disruptive side effects (44). Levodopa therapy is often the first line of pharmacologic treatment, with increasing doses often necessary to mitigate motor symptoms as PD progresses (43). Additional dopaminergic pharmacotherapies may be prescribed based on age of onset, disease severity, and side effects of levodopa monotherapy (LM) (44). Poly-DT refers to the prescription of additional DT prescribed in addition to levodopa, for example, dopamine receptor agonists, monoamine oxidase B (MAO-B) inhibitors, and Catechol-O-methyltransferase (COMT) inhibitors (45–48). Each of these medications is expected to have very different mechanisms by which the dopaminergic system is affected. Therefore, their effects and interaction with non-motor symptom co-morbidities in PD and their effect on time perception in patients may be varied.

The cause and progression of PD (progressive loss of dopamine terminals over time), associated non-motor symptoms (particularly those affecting the dopamine system), and dopaminergic pharmacotherapies used to treat PD, suggest that patients with PD will possess differences specific to the impact of their disease on their overall dopaminergic system. Increases and decreases in the efficacy of dopamine neurotransmission (and complex combinations caused by changes in different aspects of the dopaminergic system) ought to have predictable effects on dopamine-dependent processes like interval timing. Thus, we hypothesized that a multivariate approach to characterizing individual differences in patients with PD would reveal systematic differences in interval timing behavior according to dimensions of their clinical state. We tested this hypothesis in patients with PD presenting with a heterogeneous profile of comorbid symptoms (Table 1).

**Table 1:** Summary of participant demographics and clinical features.

|  | **<u>PD Patients</u>**<br>**<u>(n=19):</u>** |
| --- | --- |
| **<u>Mean Age (Years):</u>** | 65.52 ± 6.5 |
| **<u>Sex</u>** |  |
| Male: | 11 (64.7%) |
| Female: | 8 (42.1%) |
| **<u>Race</u>** |  |
| Asian: | 0 (0.0%) |
| Black or African American: | 0 (0.0%) |
| White: | 19 (100.0%) |
| 2 or More Races: | 0 (0.0%) |
| **<u>Handedness</u>** |  |
| Right: | 18 (94.7%) |
| Left: | 1 (5.3%) |
| Ambidextrous: | 0 (0.0%) |
| **<u>Clinical Features</u>** |  |
| Diagnosed with Depression: | 4 (21.1%) |
| Mean PD Duration (Years): | 4.368 ± 2.1 |
| Mean LEDD (mg): | 564.6 ± 210 |
| Mean UPDRS Score: | 19.71 ± 4.9 |
| Diagnosed with ICD: | 11 (57.9%) |
| Mean QUIP Score: | 13.00 ± 11 |
| Prescribed Poly-Dopaminergic Therapy: | 11 (57.9%) |

*PD=Parkinson's Disease; LEDD=Levodopa Equivalency Daily Dose; UPDRS=United Parkinson's Disease Rating Scale; ICD=Impulse Control Disorder; QUIP=Questionnaire for Impulsive-Compulsive Disorders in Parkinson's Disease. Data are expressed as Mean ± SD or Frequency (Percentage).*

Patients with PD performed a temporal bisection task both on and off their standard of care prescription DT (Fig. 1A) and individual differences in patients’ clinical profiles were recorded. Multiple linear regression models were fit to determine a connection between patients’ clinical profiles and psychometric measures of interval timing (Fig. 1B-C). These data revealed a clear association between interval timing and individual differences in patients’ clinical profiles, demonstrating a link between the dopaminergic mechanisms of interval timing and altered dopamine function resulting in PD symptomology. The data was then fit with a leave-one-out cross-validated principal component regression model using the psychometric measures of interval timing (Fig. 2) as independent variables to predict the specific clinical features and comorbidities associated with individual patients with PD. Our results support our hypothesis and demonstrate a clear predictable association between complex PD clinical symptomology and quantitative differences in interval timing consistent with the hypothesized roles for dopamine in both the clinical features and timing behavior. Our results suggest that relatively simple psychophysical tasks that measure interval timing may be used as a behavioral biomarker for stratifying heterogeneous PD pathology. Further, such a paradigm may be used to investigate the neurobiological mechanisms that link the dopaminergic system, time perception and motor and non-motor PD pathology.

**Fig. 1.**
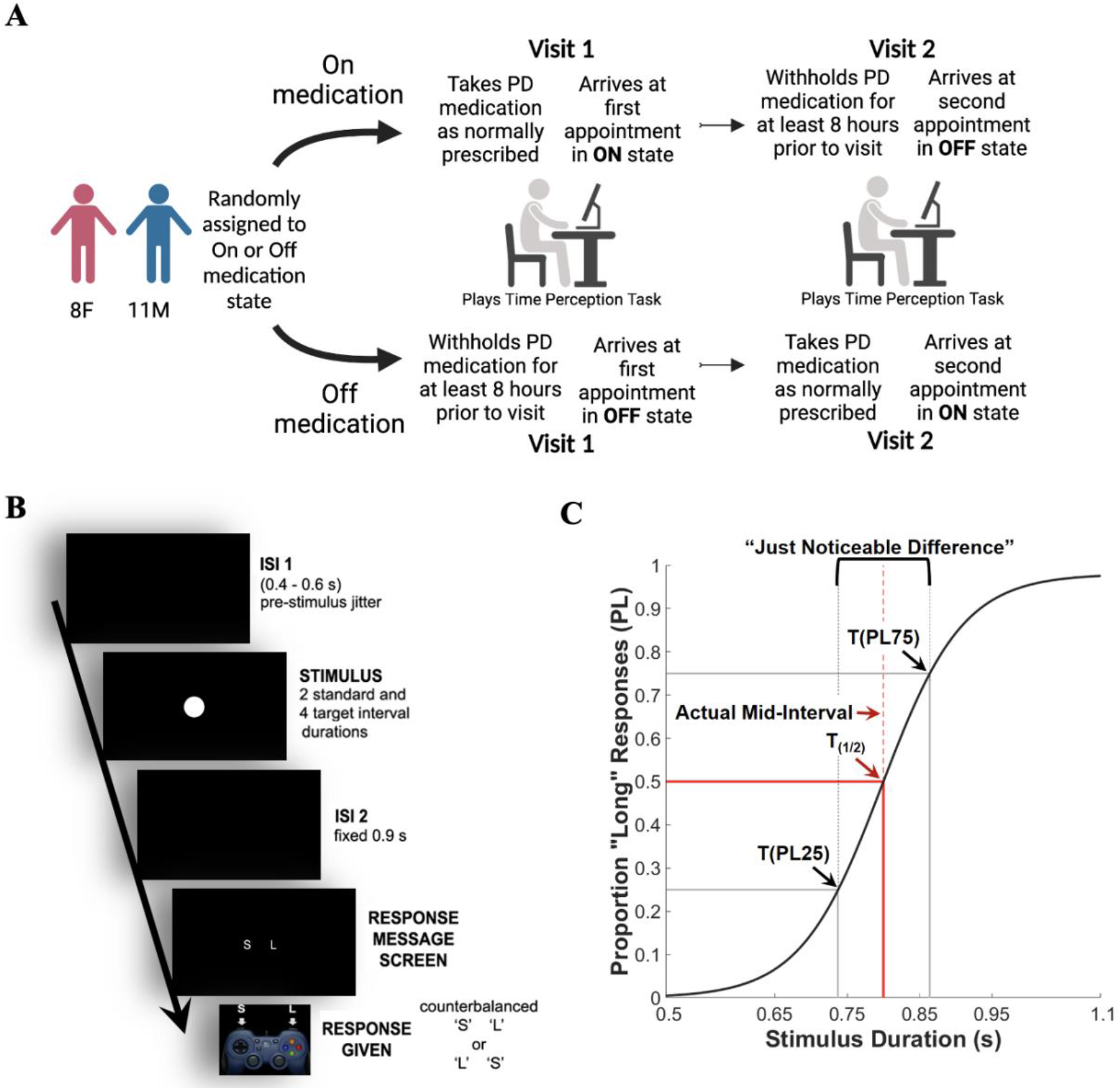
The temporal bisection task. **A.** PD patient recruitment and medication state assignment. Each patient was randomly assigned to either a first on-medication or off-medication visit. The second visit was the alternate medication state. During both visits, each patient played a temporal bisection task (Created with BioRender.com). **B.** The schedule of a single trial from the bisection task. A trial begins with an Inter-Stimulus Interval (ISI) consisting of a pre-stimulus jitter of a black screen for a poison distribution of 0.4s to 0.6s. The stimulus of a white circle is presented in the center of the screen for one of six stimulus durations: 0.5s, 0.65s, 0.75s, 0.85s, 0.95s, 1.1s. ISI 2 followed for a fixed 0.9s. The response screen then appears with a counterbalanced S and L presented on the screen. The subject judges whether the stimulus interval was closer in duration to a previously learned short (0.5s) or long (1.1s) interval and submits their response using the Logitech Controller. **C.** A psychometric function applied to data in a healthy control demonstrates near-optimal temporal bisection task performance. Psychometric indices were derived from the psychometric function, with the solid red line showing the subject’s Bisection Point (BP or T(1/2)), which is the stimulus duration at which 50% of responses were “long”. This is compared against the dotted red line of the actual mid-interval (0.8s). The difference between the BP and the actual mid-interval dictates the bisection point error (BPE). The stimulus duration corresponding to proportion long 25 (T(PL25)) refers to the duration at which a subject responded long 25% of the time whereas the duration corresponding to proportion long 75 (T(PL75)) refers to the duration at which a subject responded long 75% of the time. The difference between these values is the “just noticeable difference”. This difference divided by two yields an estimate of temporal precision, the difference limen (DL). The Weber Fraction (WF) is then calculated as the quotient of the DL and BP. For both the DL and WF, more precise performance is represented by lower values.

**Figure 2:**
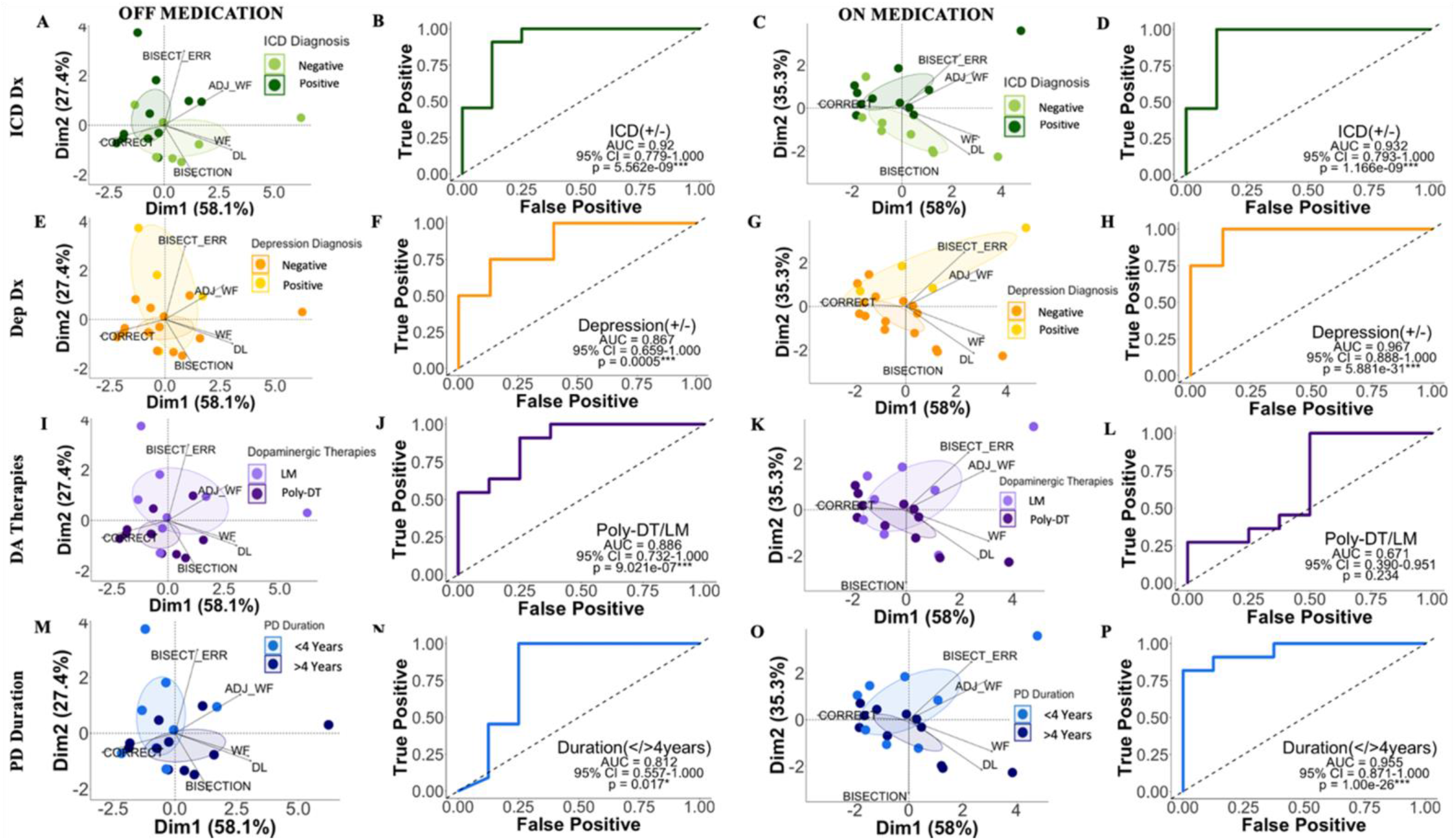
Principal Component Analyses (PCA) of the time perception metrics and receiver operating characteristic (ROC) curves illustrating the diagnostic accuracy of dimension loadings OFF and ON medication states in the discrimination of clinical subgroups of PD patients. **A.** Biplots showing loadings of dimensions one and two with 95% confidence ellipses showing the groupings of ICD(+) and ICD(-) patients off-medication were generated. **B.** ROC curve showing comparison of true and false positives for ICD diagnosis in patients with PD off-medication, with associated Area Under the Curve (AUC), 95% Confidence Interval (CI), and p-values (AUC=0.92, CI=O.78-1.0, p=5.6e-09***). **C-D.** For ICD diagnosis on-medication (AUC=0.93, CI=0.79-1.0, p=l.7eO9***). **E-F.** For depression diagnosis off-medication (AUC=0.87, CI=O.66-1.0, p=0.0005***). G-H. For depression diagnosis on-medication (AUC=0.97, CI=O.89-1.0, p=5.9e-31***). I-J. For Dopaminergic Therapies off-medication (AUC=0.89, CI=O.74-1.0, p=9.0e-07***). K-L. For Dopaminergic Therapies on-medication (AUC=0.67, CI=O.39-0.95, p=0.23). M-N. For disease duration off-medication (AUC=0.81, CI=O.56-1.0, p=0.017*). O-P. For disease duration on-medication (AUC=0.96, CI=O.87-1.0, p=1.0e-26***). *Dim=Dimension; ICD DX=Impulse Control Disorder Diagnosis; DEP=Depression; PD=Parkinson’s disease. L Levodopa Monotherapy; DT=Dopaminergic Therapies. Significance based on p<0.05*;0.01**;0.001\**

## RESULTS

### No differences in average interval timing between patients with PD on or off-medication

Patients with PD (N=19, Table 1), while on and off their DT (Fig. 1A), performed a temporal bisection task with intervals ranging from 500 to 1100ms (Fig. 1B-C). The average accuracy and precision of interval timing within participant groups were compared across medication states. Accuracy measures included the bisection point, bisection point error, and percent of trials correctly selected as long or short. Precision measures included weber fraction, difference limen, and adjusted weber fraction (please see methods section for description of each psychophysical timing metric). On average, patients with PD did not show significant differences in interval timing measures when on vs. off DT (*bisection point: p=0.63, g=0.15, β=0.35; weber fraction: p=0.80, g=0.08, β=0.32; difference limen p=0.74, g=0.11, β=0.33; percent correct: p=0.85, g=-0.06, β=0.32; bisection error: p=0.30, g=-0.34, β=0.489; adjusted weber fraction: p=0.38, g=-0.28, β=0.43)* although the Bayesian evidence was ambiguous *(>0.3 and <3.0)* (see Table S1 for all group comparisons). However, medication state did have an effect on the ability of generated predictive models to discriminate whether patients were prescribed mono-versus poly-DT (predictive models and associated results described below, and in Figure 2).

### Interval timing predicts the clinical profiles of patients with PD

For each patient with PD, profiles of clinical characteristics were collected, including comorbid diagnoses, types of DT (e.g., LM or Poly-DT; see Table S4 for prescribed DTs for each patient), Levodopa Equivalency Daily Dose (LEDD), United Parkinson’s Disease Rating Scale (UPDRS), age, PD duration, and Questionnaire for Impulsive-Compulsive Disorders in Parkinson’s Disease–Rating Scale (QUIP-RS) score (Table S2). These clinical features, including being in the on- or off-medication state, were collected to explore the heterogeneity of PD presentation in each participant, and to determine how each of these clinical features directly related to alterations in interval timing.

Six psychophysical timing measures of accuracy and precision were calculated from psychometric functions of interval timing (Figure 1C for example of a psychometric function; see supplemental methods for a detailed description of psychometric function fit). Akaike Information Criterion (AIC)-penalized (49) multiple linear regression models were used to determine associations between clinical features from patients with PD and their psychophysical timing measures. The models indicated that from the profiles of clinical characteristics, ICD (*bisection point: t=3.6, p=0.003**)* and depression diagnoses *(bisection point error: t=3.4, p=0.004**; percent correct: t=-3.3, p=0.005**; adjusted weber fraction: t=2.4, p=0.028*),* disease duration (*percent correct: t=-4.0, p=0.001**; difference limen: t=3.2, p=0.007**; weber fraction: t=3.0, p=0.01*; adjusted weber fraction: t=2.4, p=0.028*)* and multitude of DT *(bisection point: t=3.8, p=0.002**; percent correct: t=3.4, p=0.005**; difference limen: t=-3.5, p=0.004**; weber fraction: t=-3.3, p=0.005**; adjusted weber fraction: t=-3.3, p=0.004**)* were significantly associated with specific psychophysical timing metrics (see Table S3 for beta coefficients of all clinical variables that survived AIC correction and associated model p-values).

We used the associations observed from the AIC linear regression models to direct the rest of our analyses. From these results, we hypothesized that interval timing would be predictive of ICD and depression diagnoses, disease duration, and multitude of DT in patients with PD. To test this hypothesis, we performed a principal component analysis of six time perception psychophysical measures (i.e., bisection point, bisection point error, percent correct, difference limen, weber fraction, adjusted weber fraction) performed separately for both the on and off-medication states (Fig. 2, see Fig. S1 for dimension loadings). To determine if patients with shared clinical features would appear clustered together, we produced biplots displaying loadings onto the first two principal components, which together accounted for 85.5% of the variance in interval timing data for the off-medication group and 93.3% for the on-medication group. Overlaid 95% confidence interval ellipses showed distinct groupings of ICD(+/-) diagnosis (Fig. 2 A-D), depression(+/-) diagnosis (Fig. 2 E-H), prescription of Poly-DT or LM (Fig. 2 I-L), and disease duration (> or <4 years) (Fig. 2 M-P).

Next, we explored whether these clinical features could be predicted based on combinations of interval timing performance measures. Specifically, we performed a leave-one-out cross-validated multivariate logistic regression analysis using the principal component loadings as six independent predictors of these clinical groupings (Table S5). These regression models were used to produce probability values for each individual patient to classify them as ICD(+) or (-), depression(+) or (-), disease duration > or < 4 years, and prescribed Poly-DT or LM. The resulting predictions revealed high accuracy rates of greater than 70% for each clinical grouping. Accompanying receiver operating characteristic (ROC) curves were plotted for each clinical grouping and associated Area Under Curve (AUC) values were calculated (Fig. 2). The ROC curves revealed acceptable *(AUC>0.70;p<0.05*)* to excellent *(AUC>0.90;p<0.01**)* diagnostic fit (50) compared to chance level (AUC=0.50) for all included clinical features (Fig. 2), with the exception of Poly-DT in the on-medication state *(AUC=0.67;p=0.234)* (Fig. 2L).

### PD presentation can be observed in psychometric functions of interval timing

Based on the collected clinical characteristics and the associations found in our predictive models, we aimed to observe the specific differences in temporal performance using individual patient psychometric functions. We first identified and separately grouped ICD(+) and ICD(-) patients according to their ICD status, and we then modeled interval timing of ICD(+) and ICD(-) patients using psychometric functions of performance on the temporal bisection task (see Fig. 1C for an example psychometric function). The resulting psychometric functions, displaying representative performance from an ICD(+) and ICD(-) individual revealed that an ICD(+) diagnosis was associated with a right shift from the mid-interval duration (0.8s) when compared to ICD(-) subjects (See Fig. 3A for off-medication group comparison). Specifically, we observed that patients with PD with an ICD(+) diagnosis had a significantly lower bisection point than patients without an ICD diagnosis, indicating ICD(+) patients overestimated the duration of time intervals in the off-medication state, and therefore were less accurate in their time perception than ICD(-) patients (Table S6).

**Fig. 3.**
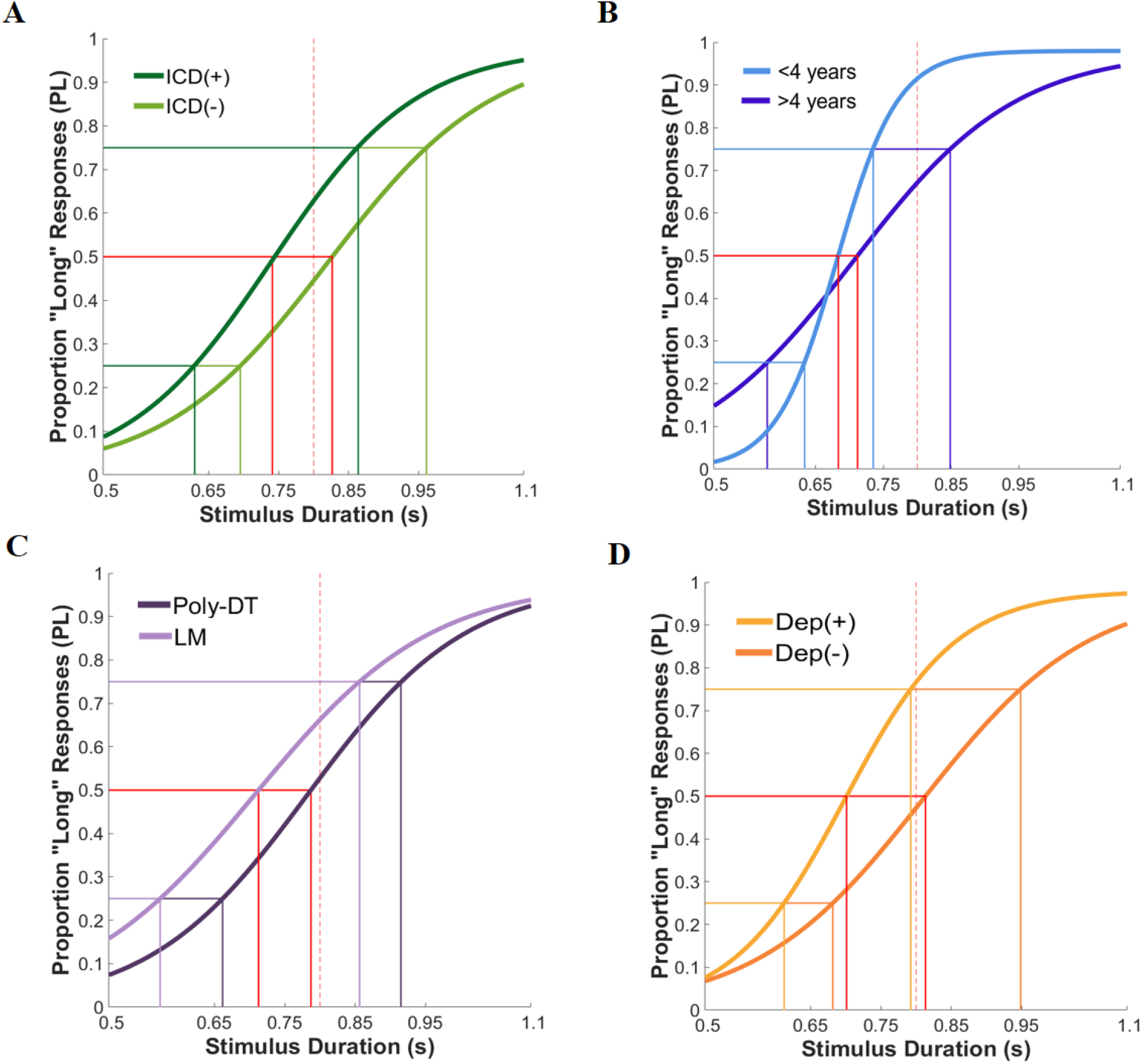
Representative psychometric functions of patients with Parkinson’s disease (PD) demonstrate the relationships between clinical features of PD and interval timing performance. **A.** Psychometric functions comparing the presentation of a representative Impulse Control Disorder (ICD)(+) patient (dark green) to the psychometric function of a representative ICD(-) patient (light green) in the off-medication state. **B.** Psychometric functions comparing the presentation of a representative patient with a PD duration of less than four years (light blue) to the psychometric function of a representative patient with a PD duration of more than four years (dark blue) in the off-medication state. **C.** Psychometric functions comparing the presentation of a representative patient prescribed dopaminergic polytherapy (dark purple) to a representative patient prescribed Levodopa monotherapy (light purple) in the off-medication state. **D.** psychometric functions comparing the presentations of a representative depression(-) patient (orange) to a representative depression(+) patient (yellow) in the on-medication state. *BP=Bisection Point; WF=Weber Fraction; PC=Percent Correct. AWF=Adjusted Weber Fraction. AIC=Akaike Information Criterion.*

Next, we considered the duration of PD for each patient, which had a group median of four years (see Table S1 for group means of clinical features and Table S4 for individual disease durations). To compare interval timing of patients with PD who had longer disease durations (≥4 years) to those who had shorter disease durations (<4 years), we directly compared the psychometric functions of interval timing between two representative patients in each group. We found that subjects with a longer disease duration had flatter psychometric functions, reflecting poorer temporal precision, compared to subjects with a shorter disease duration across both medication states (Fig. 3B).

To observe the effect of the multitude of prescribed DT on time perception, we grouped patients with PD based on their prescription of LM or Poly-DT. We displayed the psychometric functions of two representative patients from each group in the off state of medication (see Table S2 for types of DTs). The resulting psychometric functions of patients who were prescribed LM exhibited large leftward shifts from the midpoint and significantly shallower slopes than patients prescribed Poly-DT (Fig. 3C). Thus, patients prescribed Poly-DT demonstrated both better temporal precision and timing accuracy than patients prescribed LM (Fig. 3C).

Finally, we identified patients with comorbid depression diagnoses and compared the psychometric functions of one representative depression(+) patient and one depression(-) patient. The resulting psychometric functions showed that depression(+) patients exhibited a large leftward shift from the mid-interval compared to depression(-) patients (Fig. 3D). Depression(+) patients presented with a higher bisection point error, higher adjusted weber fraction, and a lower percent correct than depression(-) patients with PD, demonstrating more variability in their bisection point and overall poorer temporal precision and accuracy reflected in their individual psychometric functions.

### Individual differences in complex combinations of clinical characteristics of PD relate to interval timing

Based on the results of our PCA and multivariate regression analyses, we examined whether individual combinations of clinical features would predict individual features of psychometric functions. To investigate, we first selected Subject 3, a patient who presented with negative ICD and depression diagnoses, and was prescribed Poly-DT. These are clinical characteristics that our analyses showed to result in a slight underestimation of intervals, but overall precise timing. However, Subject 3 also had a disease duration greater than 4 years (Table S4). Therefore, with the combination of these clinical features and the associations we found in our model, we predicted Subject 3 would be less temporally precise due to a longer disease duration. This prediction was supported by Subject 3’s psychometric function, which displayed a slight rightward shift in BP close to the mid-interval and a shallower function (Fig. 4A). We then considered Subject 17, who presented with a similar clinical profile as Subject 3 – prescribed Poly-DT, ICD(-), depression(-) – but this patient had a disease duration less than 4 years. We predicted that due to their shorter disease duration, Subject 17 would have superior temporal precision than Subject 3, but otherwise similar interval timing performance. In fact, Subject 17’s psychometric function showed a rightward shift in the BP close to the mid-interval and a steeper function demonstrating more precise interval timing (Fig. 4B). Similarly to Subject 17, Subject 1 was prescribed Poly-DT, was depression(-), and had a disease duration less than 4 years, but was ICD(+). On the basis of our previous analyses, we expected that this subject would exhibit relative overestimation of time intervals due to being ICD(+). Consistent with this prediction, this subject exhibited a psychometric function with a leftward shift in their BP, and otherwise similar interval timing performance as Subject 17 (Fig. 4C). Subject 13 had a similar clinical profile to Subject 1, but was prescribed LM, was both depression(+) and ICD(+), and had a disease duration less than 4 years. Due to the concurrence of being prescribed LM and being depression(+), we expected that they would display poorer temporal accuracy than Subject 1. In support of this prediction, their psychometric function showed a large leftward shift in the BP away from the mid-interval (Fig. 4D).

**Fig. 4:**
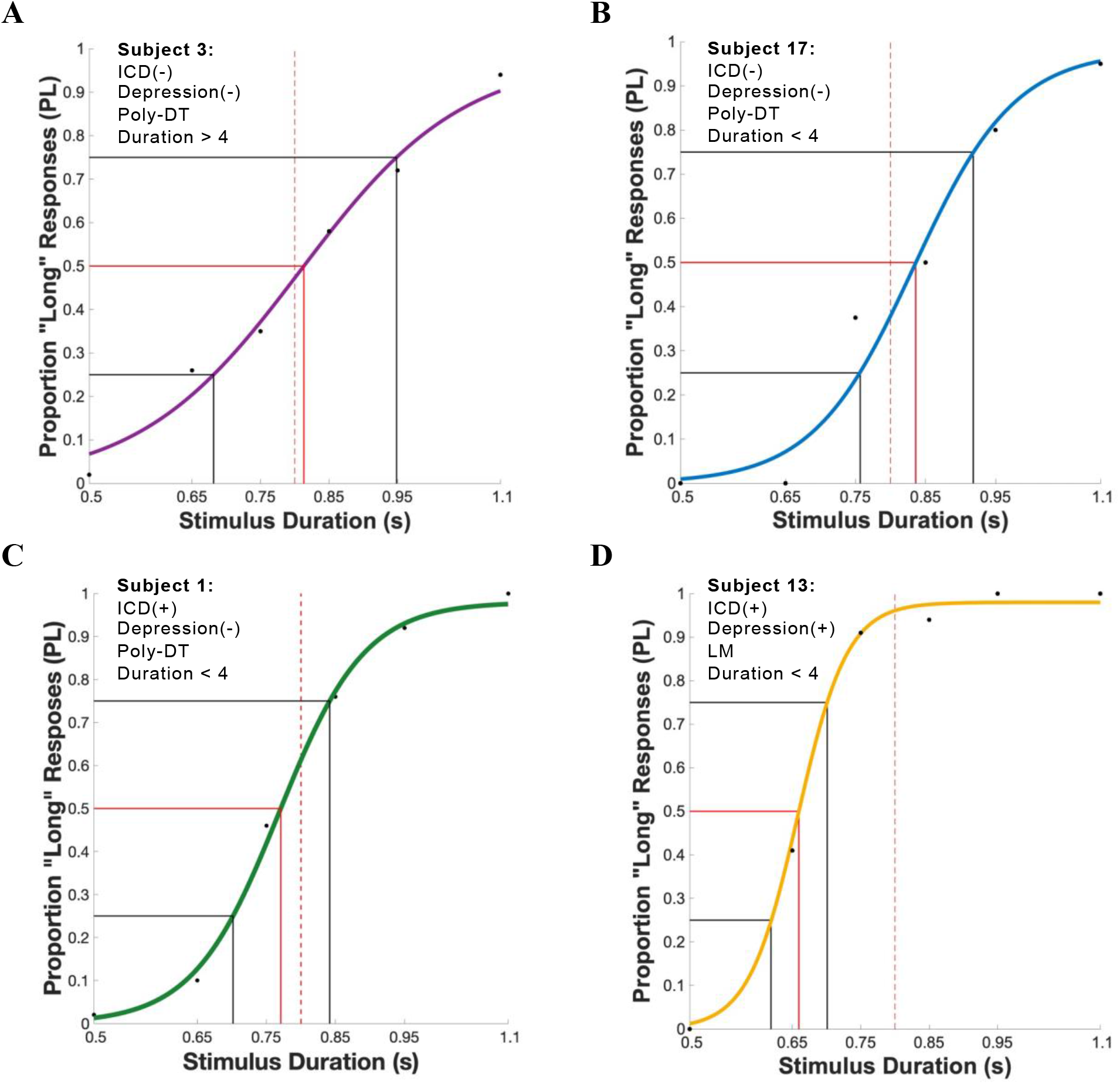
Psychometric functions demonstrate relations between multiple comorbidities and heterogeneous clinical features of Parkinson’s disease (PD) to temporal performance. **A.** The combination of clinical features of Subject 3 is represented in a psychometric function that displays relatively high accuracy, but lower precision (BP:0.818; WF:0.162; PC:73.09). **B.** In comparison to A, the psychometric function of Subject 17 shows relatively high accuracy and precision (BP:0.831; WF:0.133; PC:75.34). **C.** In comparison to B, the psychometric function of Subject 1 shows high precision, but an overestimation of time intervals (BP:0.775; WF:0.091; PC:82.17). **D.** In comparison to C, the psychometric function of Subject 13 shows high precision and low accuracy with an overestimation of time (BP:0.665; WF:0.060; PC:72.42). *DT=Dopaminergic Therapies; BP=Bisection Point; WF=Weber Fraction; PC=Percent Correct.*

Overall, patient performance on the interval timing task corroborated our model predictions, and we found that these predictions were consistent with the co-occurrence of multiple clinical features of PD. Additionally, we showed that psychometric functions of interval timing performance could be used to identify and observe individual differences in clinical presentation and comorbidities of PD (see Fig. S2 for all patient psychometric functions). Therefore, examination of psychometric functions of interval timing performance could provide insight into the comorbidities and clinical presentation of patients with PD.

## DISCUSSION

The present study investigated time perception in patients with PD with varying disease duration, variable medication regimens, and comorbid non-motor symptoms, including ICD and depression. We investigated time perception in these patients while they were on- and off-their prescribed dopaminergic medications. Accounting for their specific clinical state (e.g., specific motor and non-motor symptoms and medications used) proved to be critical in interpreting changes in their timing behavior. Performing a cross-validated, leave-one-out, multivariate regression analysis revealed systematic patterns of timing behavior that are predictive of individual-level clinical states involving putative changes in the dopaminergic system. The relative simplicity of interval timing tasks and quantitative analyses of resulting behavior suggest the potential for future development of time perception as a behavioral biomarker for patients with PD with complex clinical profiles of motor and non-motor symptoms.

Interval timing may not be the first cognitive process one thinks about in the context of PD. However, with the amassing of evidence that striatal dopamine regulates interval timing (7–18,51), that patients with PD exhibit alterations in timing behavior (22–23,52–55), and our currently presented evidence that interval timing can predict non-motor symptoms of PD, interval timing tasks may help to elucidate the dopaminergic mechanisms of the non-motor symptoms of PD. Research into time perception in patients with PD has given insight into how dopamine may affect behavior and cognition for many decades (52–55). Previous work has revealed that patients with PD withholding their prescribed dopaminergic therapies exhibit poor temporal accuracy (54,55), but that accurate timing can be restored with the reintroduction of these therapies (52). These medication state-based studies, however, often do not consider the heterogeneity of PD symptomology affecting timing performance. Therefore, other factors of PD, such as individual differences in motor and non-motor symptom presentation, could be confounding these outcomes. More recently, interval timing studies have begun to relate individual features of disease presentation to differences in temporal behavior. Specifically, Merchant et al. observed a subpopulation of patients with PD that presented with similar disease duration, UPDRS scores, and Levodopa equivalency daily dose that had difficulty perceiving subsecond intervals of time when compared to patients with PD that exhibited different clinical profiles (23). Additionally, Kent et al., showed that time perception performance can differentiate between specific features of psychiatric disorders and potentially serve as a useful tool in the differential diagnosis of psychiatric illnesses (56). In this study, we aim to merge these ideas to determine if the heterogeneous profiles of patients with PD, which includes psychiatric comorbidities, could be predicted with interval timing.

Interval timing tasks, like the temporal bisection task in this study, are relatively simple to perform and do not take long to complete (for instance, this task takes less than 30 minutes, which can potentially be shortened). Therefore, they may serve as an additive to standard of care appointments for patients with PD; however, designing the optimal time perception task (57) is an important consideration prior to implementation. Specifically, it is important to consider the duration of stimuli to be tested. Research examining the effects of the duration of stimuli tested during timing tasks shows that performance differs based on dopamine availability. Patients with PD have demonstrated impairments on suprasecond intervals of time when off of their prescribed DTs; but, medication state did not impact subsecond interval time perception the same patients (52,54–55). In our study, we show a consistent lack of effect of medication state on subsecond interval timing (54, Table S1), but we show that subsecond stimuli could predict other aspects of dopamine-specific clinical profiles of patients with PD (Fig. 2). Therefore, in designing time perception tasks to be used in a clinical setting, it is critical to select an optimal task design that relates to the symptomology of interest.

In this study, we investigated time perception behavior on a temporal bisection task in patients with PD while they were on- and off- of their prescribed dopaminergic medications. Each patient, in consultation with their clinical provider, develops a tailored medication regimen that aims to ameliorate their PD symptoms while also minimizing unwanted side-effects (44). Our results show that patients with PD on their prescribed DT did not differ in their interval timing compared to patients off their prescribed DT (Table S1). As studies have shown that the effects of dopamine medications differ based on stimulus duration (52,54–55), our results corroborate a finding that withholding DT does not alter time perception in the subsecond duration range (54). This result was internally validated across on and off-medication states for each comorbidity (Fig. 2). A possible explanation for this lack of observed difference could be that we recruited a heterogenous population of patients with additional psychiatric comorbidities, whereas other studies often exclude these patients to compare more homogeneous groups of patients with similar disease profiles. Therefore, the complex combination of neurological and psychiatric disorders, as well as individualized medication profiles, could be masking the on-versus off-medication effects seen in past studies. More research is needed to determine the circuit and receptor-specific interactions potentially modulating this lack of medication effect, but altogether this highlights the complexities of PD populations that simply considering these patients as a homogenous “movement disorder” cohort undermines.

Due to heterogeneity in PD symptoms, patients are often prescribed multiple therapies that differentially target their dopamine systems to treat their symptoms (44). Therefore, we explored the impact of patients prescribed solely Levodopa monotherapy (LM) compared to a regime of Poly-DT. Our results reveal that patients prescribed Poly-DT are both more precise and more accurate in their interval timing than patients prescribed LM, across on and off-medication states (Table S1). Additionally, we found that patients prescribed LM versus Poly-DT could be predicted based on interval timing performance in the off-medication state; however, the predictability of prescription of Poly-DT disappeared when patients were on their PD medications (Figure 2). This result could stem from clinicians avoiding the prescription of dopamine agonists and certain poly-DT to patients vulnerable to psychiatric comorbidities, including ICD. Therefore, patients prescribed poly-DT may have been cognitively more robust at baseline resulting in better timing performance. Future research into the effects of DT on interval timing would need to include cognitive measures, like the Montreal Cognitive Assessment, to control for cognition at baseline. More research could also shed light on the underlying dopaminergic mechanisms of the non-motor effects of dopaminergic medications in patients with PD, as well as aid in the medication management of patients with PD.

Patients with PD present in the clinic with motor symptoms at the core of their diagnosis; however, each patient presents with a range of severity in motor and non-motor symptoms that may be independent of, caused by, or exacerbated by their dopamine therapies (58). For example, ICD is a behavioral addiction believed to be induced by the prescription of dopamine receptor agonists in patients with a preferential affinity for dopamine (D3) receptors in the striatum (27–32). The preferential interaction between dopamine receptors and dopamine medications is believed to underly the onset of aberrant behaviors in the form of compulsive and repetitive gambling, sex, buying, and/or eating (59–61). Other behavioral ramifications of comorbid ICD in patients with PD, such as timing behaviors, were previously unknown. However, investigations into impulsivity in populations outside of patients with PD, such as schizophrenia and borderline personality disorder, demonstrate an association between impulsive behaviors and increased accuracy and precision variability in interval timing (38–40). In this study, we found that ICD(+) patients with PD tend to overestimate intervals of time compared to ICD(-) patients (Figure 3A), which is consistent with previous studies on impulsivity in timing (39–40). As pharmacological studies have shown that dopamine agonists tend to yield a relative leftward shift in psychometric functions (8,11), it has been hypothesized that increased levels of dopamine in the striatum are associated with the overestimation of time intervals (51). Therefore, as the interaction of overactive dopamine receptors with dopamine agonists are thought to produce a state analogous to a hyperdopaminergic state in patients with ICD (31,32), our results align with the hypothesized role of dopaminergic dysfunction in patients with PD with comorbid ICD.

Another common comorbidity of PD is depression (33,34), of which the etiology remains unclear (58). The extant literature, however, supports the depletion of dopaminergic tone in the basal ganglia in depressive patients (58,59). This is consistent with the action of many antidepressants, which aim to increase dopamine by targeting the midbrain (62,63). Decreased dopamine activity has been linked to decreased temporal precision (2,6), and patients with depression have been shown to exhibit a chronic underestimation of time (41,42). Our results show that depression in patients with PD may not be as congruous. We found significant variability in timing accuracy with some patients overestimating intervals and some patients underestimating intervals of time (Table S1). This outcome could relate to individual differences in depression diagnosis and prescribed medications used to treat depression symptoms, which patients were not asked to withhold. It could also provide insight into how depressive symptoms in patients with PD differ from other forms of depression. Future time perception research into comorbid depression in patients with PD could help explain this significant variance in timing accuracy and help determine the underlying role of dopamine in depression.

In this study, we showed that performing a cross-validated, leave-one-out, multivariable regression analysis can reveal systematic patterns in individual-level timing behavior that are predictive of individual-level clinical states involving the putative changes in the dopaminergic system. The results of our study demonstrate that behavior on a simple time perception task can predict ICD diagnosis, depression diagnosis, disease duration, and multitude of dopamine medications in patients with PD with high diagnostic accuracy (Figure 2). These findings expand upon previous work on time perception differences based on individual differences in PD severity (64), and it shows that beyond just correlation, interval timing can be predictive of individual clinical features of PD. A limitation of our study is the small sample size of patients with PD (n=19). Future work investigating time perception would need to be completed using a larger sample of patients with PD and involve the specific recruitment of patients with comorbidities. It would also be prudent to utilize a more diverse sample of patients, with more severe disease progression and duration. Another limitation is the utilization of a temporal bisection task design, which presents subjects with a brief training phase followed by a testing phase, with no feedback on performance during either phase. This task design presents with many idiosyncrasies in human performance that cannot be fully explained by currently proposed models of time perception (57). Additionally, significant differences in performance have been noted between animals and humans (57). Therefore, care must be taken when interpreting results from temporal bisection tasks and future work is still needed to determine associated timing mechanisms.

More work is needed, but basic cognitive neuroscience has a variety of tasks that aim to probe neuromodulatory systems (especially the dopamine system). The relative simplicity of interval timing tasks and quantitative analyses of resulting behavior suggest the potential for future development of time perception behavior as a behavioral biomarker for patients with PD with complex clinical profiles that include motor and non-motor symptoms (65,66). The need for accurate biomarkers of PD has been proposed, with the objective of producing biomarkers predictive of disease progression (66,67). Our study shows that with future development, time perception has the potential to be used as a behavioral biomarker for individual differences in progressive dopaminergic dysfunction associated with PD.

## MATERIALS AND METHODS

### Subjects

Patients with PD (n=19) were recruited through Atrium Health at Wake Forest Baptist’s (AHWFB) movement disorders clinic (Table 1). All patients with PD were phone-screened by a researcher to determine eligibility prior to any scheduled research visits. Patients were eligible to participate if they were between the ages of 21-85 years, had a confirmed diagnosis of PD from a trained movement disorder specialist, were currently prescribed and taking DT, and had the ability to withhold DT for up to 11 hours (8 hours prior to experiment and 2-3 hours for the duration of testing). Patients with PD were ineligible if they failed to meet the inclusion criteria, had moderate or severe dementia, or the inability to use a computer. All patients provided informed written consent in accordance with approval by the IRB committees at AHWFB (IRB00017138).

### Study design

The study took place over two visits, a minimum of one week apart (Fig. 1A). During one visit, patients were asked to withhold their DT prescribed to treat their PD symptoms for at least 8 hours prior to the visit (“off-state”). During the other visit, patients were asked to take DT as normally prescribed (“on-state”). Medication state was randomized for the first visit via a coin toss and followed by a second visit of alternate medication state. Clinical measures were collected for each patient, including age, PD duration, number and type of DT, and comorbid diagnoses (Table 1). Clinical data was collected from a combination of self-report surveys and clinical records of patients utilizing EPIC software from AHWFB (Table S2). ICD diagnosis was measured by the QUIP-RS utilizing diagnostic thresholds from Weintraub et al. 2012 (68). The QUIP-RS was completed by each PD patient at the end of their second research visit. Depression diagnosis was measured by physicians and reported through clinical records and/or by self-report questionnaires. Disease motor severity was measured by UPDRS reported through clinical records of neurology visits at AHWFB (69). Levodopa equivalency daily dose was calculated from reported prescribed DT utilizing dopaminergic equivalency conversions from EQUIDopa platform (70) (Table S4).

### Visual display and computer set-up

The task was performed in Matlab R2020b, using Psychophysics Toolbox extensions (71). Subjects were seated about two feet from a computer screen and used a Logitech gaming controller to ledge their judgments. During the task, an image of a white circle was shown on a black computer screen centered at 2-4° of their visual field. Subjects submitted their responses using the shoulder keys on the Logitech controller.

### Temporal bisection task

The task was broken up into two phases, a training and a testing phase. The participant first learnt two anchor time intervals, a short interval of 0.5s and a long interval of 1.1s. A total of ten short anchor intervals and ten long anchor intervals appeared randomly during the training phase. If the participant did not score greater than sixty percent correct during the training phase, they were automatically presented with additional trials until the accuracy threshold was reached or ten minutes had passed. The task would automatically move onto the testing phase after ten minutes irrespective of the accuracy threshold. Six patients needed to repeat the training phase of the task, and two were unable to meet the accuracy threshold before moving onto the testing phase. Data from the training phase was not used in the above analyses. The testing phase included additional target intervals of 0.65s, 0.75s, 0.85s, 0.95s, as well as the 0.5s and 1.1s anchor intervals. The participant judged whether the presented stimulus intervals were closer in duration to the short or long anchor intervals learned during the training phase. The three shortest intervals — 0.5, 0.65, and 0.75 seconds — were counted as correct if the subject responded with short, while the three longest intervals — 0.85, 0.95, and 1.1 seconds — were counted as correct if the subject responded with long.

Each trial in the task (Fig. 1B) began with a black screen that lasted for 0.4-0.6s (inter-stimulus interval one, ISI 1). The ISI 1 was drawn randomly from a truncated Poisson distribution. It was immediately followed by a white circle in the center of the black screen (displayed for 0.5s, 0.65s, 0.75s, 0.85s, 0.95s, and 1.1s), and a fixed 0.9s interval of black screen (inter-stimulus interval two, ISI 2). Subsequently, a response screen appeared, with either the “L” and “S” or the “S” and “L” letters shown for the participant to indicate whether the stimulus was perceived as closer in duration to the short or long anchor (“S” for “short” and “L” for “long”). Subjects submitted their response by pressing a button on the side of the controller that corresponded with that of a response letter on the screen. To address motor and attentional biases that are unable to be controlled for in rodent-based studies, the order of response letters on the screen was random. At the completion of each block of trials, a screen would appear to signify the start of a new block.

The task was performed in two sessions. Both sessions comprised the training and testing phases. However, they both differed in number of testing-phase trials. There were four and six blocks of fifty trials each in the first and second session, respectively. Each anchor appeared five times and each target interval ten times in a random order within each block, totalling to 200 and 300 trials in the first and second session, respectively. Both sessions (500 total trials) were played during each medication state visit about a week apart.

### Psychometric measures

We generated psychometric functions of temporal behavior by fitting logistic curves to the proportion of “long” responses across stimulus intervals using Palamedes Toolbox version 1.11.2 in Matlab R2020b (Fig. 1C; see supplemental methods for parameter values and fitting procedure) (72–77). From these psychometric functions, psychophysical indices for temporal accuracy and precision are calculated. Temporal accuracy is measured by the bisection point (BP), the duration at which 50% of participant responses were long, and bisection point error (BPE) measured by (72):

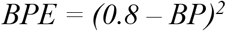

Temporal precision was measured by the difference limen (DL), weber fraction (WF), and adjusted WF (73,74).

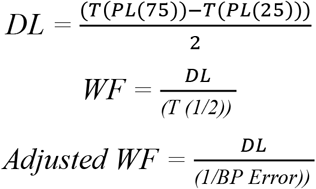

The percentage of trials each participant correctly categorized as long or short was calculated as an additional accuracy measurement. Mean psychometric measures were compared at a group level between patients with PD in the on- and off-medication states (Table 1) (76).

### Stepwise AIC linear regression models

To determine if there were associations between time perception behavior and clinical features of PD, time perception performance metrics (bisection point, bisection point error, percent correct, weber fraction, adjusted weber fraction, difference limen) were used as dependent variables in a stepwise linear regression analysis using bidirectional elimination to determine the combination of predictor clinical measures (medication state, age, disease duration, multiple medications, levodopa equivalency daily dose, depression diagnosis, ICD diagnosis, QUIP score, UPDRS score) whose inclusion generated the lowest AIC score (Table S2). AIC linear regression models were performed on time perception performance metrics collected and analyzed at the group level, from patients with PD in both the on (n = 19) and off (n = 19) medication states, and all patients with PD pooled together (n = 38). Regression results were considered significant for p < 0.05.

### Leave-one-out cross-validated multivariate logistic regression model

Time perception performance metrics (bisection point, bisection point error, percent correct, weber fraction, adjusted weber fraction, difference limen) were utilized in a PCA, resulting in six principal components (see Fig. S1 for PCA results, Fig. 2 for biplots of principal components one and two). These principal components were used as independent variables in a leave-one-out cross-validated multivariate logistic regression model using clinical features found from the initial AIC linear regression analyses. From this model, the probability of each patient with PD presenting as ICD(+), depression(+), having a PD duration >4 years, and prescribed poly-DT was estimated. If the resulting probability was greater than 0.5, then we predicted that patient with PD would possess that clinical feature. From these predictions, the number of true and false positives were accumulated, and ROC curves were generated to show the diagnostic accuracy of timing behavior for each clinical feature (Fig. 2). Diagnostic accuracy was measured via Delong’s method (50) with the calculation of AUC, Confidence Interval (CI), and difference of AUC value of predictive model from chance (chance AUC = 0.5). Calculations and modeling were generated via Palamedes Toolbox in Matlab and Rstudio Statistical Software (76,77).

## Acknowledgments

We would like to acknowledge Mary Moya-Mendez for her contributions to the IRB writing process.

## Funding

National Institutes of Health grant KL2TR00142(KTK)

National Institutes of Health grant R01 DA048096(KTK)

National Institutes of Health grant R01 MH121099(KTK)

National Institutes of Health grant R01 NS092701(KTK)

Biotechnology and Biological Sciences Research Council BB R01583X 1(RS,DBT)

## Author contributions

Conceptualization: ED, BL, IUH, MSS, KTK

Methodology: ED, RS, AJ, BL, DBT, KTK

Investigation: ED, AJ, BL, IUH, MSS

Visualization: ED, RS, REJ, KTK

Funding Acquisition: DBT, KTK

Project Administration: ED, DBT, KTK

Supervision: ED, IUH, MSS, DBT, KTK

Writing–original draft: ED, KTK

Writing–review & editing: ED, RS, AJ, REJ, BL, IUH, MSS, DBT, KTK

## Competing interests

Authors declare they have no competing interests.

## Data and materials availability

All data are available in the main text or supplementary materials.

## Supplementary materials

1. Table S1: Mean PD group performance measures on the temporal bisection task.

2. Table S2: Clinical variables for all PD patients that were utilized as independent variables for the AIC linear regression model.

1. Table S3: Dependent variables for linear regression analysis and the independent variables that were included (beta coefficient) or excluded(-) from the AIC generated linear regression models.

2. Table S4: Doses and types of dopaminergic therapies of all PD subjects.

3. Table S5: Beta coefficients from multivariate logistic regression models utilizing dimensions from principal component analysis as independent predictors of dependent clinical features of PD presentation for both on and off-medication groups.

4. Table S6: Results of Fischer Exact Tests to compare the categorical independent variable, impulse control disorder, for PD patients off-medication based on the dependent variable, the bisection point.

5. Figure S1: Dimension contributions from principal component analysis utilizing time perception measures for both on- and off-medication groups.

6. Figure S2: Psychometric functions of interval timing from all 19 patients with PD.

## Notes

### Competing Interest Statement

The authors have declared no competing interest.

