## supplemental_information for "Time perception reflects individual differences in motor and non-motor symptoms of Parkinson’s disease"

**Supplementary Materials**:

### Supplementary methods:

*Psychophysical Timing Metrics:* Utilizing the Palamedes Toolbox version 1.11.2 in Matlab R2020b (76), psychometric functions were fit to individual participant response patterns during the temporal bisection task. The psychometric function model fit was tested with a bootstrapping procedure in the Palamedes toolbox which involved computing the deviance from fitted values for simulated datasets (1,000 iterations). Alpha (threshold; stimlevels(2,1):.01:stimlevels(end -1,1)) and beta (slope; 0.5:.05:10) parameters were selected (76). Parameters for gamma (0) and lambda (0.02) were fixed (76). The goodness of fit p-value (pDev) was subsequently quantified as a proportion of simulated deviances larger than the deviance in the experimental data, with poorly fit functions showing pDev smaller than 0.03. No functions were eliminated due to poor fit. The psychometric parameters Bisection Point (BP), Difference Limen (DL), and Weber Fraction (WF) were calculated for each control participant and patient in each treatment state. The BP ($T_{(1/2)}$) was defined as the duration where short responses and long responses occurred with equal frequency; thus, it was the duration that produced 50% long (PL) responses. Overestimation of stimulus intervals is reflected in a BP lower than the mid-interval whereas underestimation of is reflected in a higher BP (69). In this task, the optimal BP would have occurred at 0.8s (Mid-Interval), as it was the value half-way between the long and the short interval durations. DL and WF served as two measurements of temporal precision. The DL was defined as the smallest change in the stimulus that resulted in a substantial behavioral change, otherwise known as the "just noticeable difference" divided by two. The DL was inversely proportional to the slope of the psychometric function; thus, the steeper the slope, the smaller the DL. In the case of DL, the lower the value, the more precise the performance. DL was measured by finding the difference between from values, $T\big(PL(75)\big)$, which was the time at which the respondent answered 75% PL, and $T(PL(25))$ was the time at which the respondent answered 25% PL. This value was then divided by two to calculate the DL. The WF stems from Weber's Law, which seeks to establish a linear relationship between perceptual accuracy and stimulus magnitude. A low WF was indicative of high discrimination between changes in stimulus; likewise, a high WF indicated low discrimination between changes in stimulus. A lower WF reflects a psychometric function that appeared step-like, whereas a high WF reflects a psychometric function that appeared gradual (17).

*Post-hoc Testing:* Based on BP, DL, and WF, independent t-tests were used to detect differences between and across the groupings, named "Healthy" for the control, "PD ON" for Patients with PD performing the task while taking medication, and "PD OFF" for Patients with PD performing the task while withholding their medication (Table 1). To determine effect size, the Hedge's g statistic was calculated (78). To determine the relationship between ICD and estimation bias, Fischer Exact Tests were used to calculate the odds ratios comparing the number of ICD positive and negative patients' tendencies to over or underestimate long intervals (Table S3).

*Additional Discussion of Results:* The effects of age progression on timing are widely reported in time perception literature (79,80). In our models, age was not found to be significantly associated with temporal performance ($p > 0.05$) (Supp. Table 1). Through additional analysis, the age of patients with PD was not significantly associated with their disease duration ($p > 0.05$), supporting our proposal that timing deficits seen in PD disease progression are not due to aging effects.

Table S1: Mean Parkinson's disease group performance measures on the temporal bisection task.

| | BP | P-value | g | *Bf* | WF | P-value | g | *Bf* | DL | P-value | g | *Bf* | PC | P-value | g | *Bf* | BP Err | P-value | g | *Bf* | Adj. WF | P-value | g | *Bf* |
|---|---|---|---|---|---|---|---|---|---|---|---|---|---|---|---|---|---|---|---|---|---|---|---|---|
| **OFF** | 0.769 ± 0.061 | 0.637 | 0.151 | 0.345 ± 0.000 | 0.143 ± 0.052 | 0.799 | 0.081 | 0.323 ± 0.000 | 0.111 ± 0.046 | 0.741 | 0.106 | 0.329 ± 0.000 | 74.5 ± 5.04 | 0.849 | -0.061 | 0.320 ± 0.000 | 0.005 ± 0.005 | 0.301 | -0.335 | 0.489 ± 0.000 | 0.0005 ± 0.0007 | 0.384 | -0.280 | 0.428 ± 0.000 |
| **ON** | 0.758 ± 0.072 | | | | 0.139 ± 0.047 | | | | 0.107 ± 0.041 | | | | 75.0 ± 5.70 | | | | 0.007 ± 0.009 | | | | 0.0008 ± 0.0011 | | | |

*Independent two-group t-tests. g refers to Hedge's G post-hoc tests. Bf refers to BayesFactor; expressed as percentage. Data are expressed as Mean ±SD. PD = Parkinson's disease; BP = Bisection Point; Adj. = Adjusted; Err = Error; WF = Weber Fraction; DL = Difference Limen; PC = Percent Correct. Signficance based on P-value<0.05.*

Table S2: Clinical variables for all Parkinson’s disease (PD) patients that were utilized as predictor variables for the Akaike Information Criterion (AIC) linear regression model.

| ID | Age (Years) | Disease Duration (Years) | LEDD (mg) | UPDRS Score | Depression Diagnosis | ICD Diagnosis | QUIP Score | Multiple DTs |
|---|---|---|---|---|---|---|---|---|
| 1 | 65 | 3 | 600 | 23 | - | + | 11 | + |
| 2 | 65 | 7 | 470 | 19 | - | - | 7 | + |
| 3 | 62 | 5 | 700 | 29 | - | - | 5 | + |
| 4 | 76 | 6 | 733 | 20 | - | - | 2 | + |
| 5 | 77 | 1 | 300 | 17 | - | - | 4 | - |
| 6 | 60 | 2 | 600 | n/a | + | + | 19 | - |
| 7 | 65 | 5 | 415 | 14 | - | + | 34 | + |
| 8 | 61 | 2 | 300 | 13 | - | - | 2 | - |
| 9 | 78 | 6 | 1000 | 19 | - | - | 4 | - |
| 10 | 67 | 7 | 610 | n/a | - | + | 11 | + |
| 11 | 63 | 6 | 400 | 22 | - | + | 36 | + |
| 12 | 60 | 3 | 750 | 23 | - | + | 15 | - |
| 13 | 51 | 2 | 450 | 19 | + | + | 31 | - |
| 14 | 65 | 5 | 950 | 12 | + | + | 21 | - |
| 15 | 67 | 5 | 650 | n/a | - | + | 17 | + |
| 16 | 67 | 8 | 350 | 27 | + | - | 4 | + |
| 17 | 65 | 1 | 300 | 23 | - | - | 0 | - |
| 18 | 70 | 5 | 450 | 19 | - | + | 12 | + |
| 19 | 61 | 4 | 600 | 23 | - | + | 12 | + |

*(+) = Positive Diagnosis;(-) = Negative Diagnosis; LEDD = Levodopa Equivalency Daily Dose; UPDRS = United Parkinson’s Disease Rating Scale; ICD = Impulse Control Disorder; QUIP = Questionnaire for Impulsive-Compulsive Disorders in Parkinson's Disease; DT = Dopaminergic Therapies.*

Table S3: Dependent variables for linear regression analysis and the independent variables that were included (beta coefficient) or excluded(-) from the Akaike Information Criterion (AIC) generated linear regression models.

| Group | Dependent Variable | Independent Variables in AIC Generated Model | | | | | | | | | F-stat | p-value |
|---|---|---|---|---|---|---|---|---|---|---|---|---|
| | | State | Age | Dis Dur | Mult Meds | LEDD | Dep Dx | ICD Dx | QUIP | UPDRS | | |
| PD ON | **BP:** | n/a | 0.011 | - | 0.100 | 0.020 | - | -0.087 | - | -0.035 | 9.685 | 4.95e-4*** |
| | **DL:** | n/a | - | 0.034 | - | - | 0.030 | -0.041 | - | -0.012 | 6.533 | 0.003** |
| | **WF:** | n/a | - | 0.050 | -0.063 | -0.012 | - | - | - | - | 5.321 | 0.011* |
| | **PC:** | n/a | - | -3.766 | - | - | -9.034 | 4.470 | - | 2.014 | 4.093 | 0.021* |
| | **BP Error:** | n/a | - | - | - | - | 0.013 | - | - | - | 11.31 | 0.004** |
| | **Adj. WF:** | n/a | - | - | - | - | 0.001 | - | - | - | 5.734 | 0.028* |
| PD OFF | **BP:** | n/a | 0.018 | - | 0.031 | 0.025 | - | -0.069 | - | - | 10.91 | 3.10e-4** |
| | **DL:** | n/a | 0.015 | 0.032 | -0.073 | - | - | - | - | 0.017 | 6.689 | 0.003** |
| | **WF:** | n/a | 0.012 | 0.037 | -0.084 | - | - | - | - | 0.021 | 5.261 | 0.008** |
| | **PC:** | n/a | - | -3.741 | 7.293 | - | -3.764 | 3.117 | - | -0.456 | 5.291 | 0.007** |
| | **BP Error:** | n/a | - | - | - | 0.002 | 0.005 | 0.004 | 0.002 | -0.002 | 3.487 | 0.032* |
| | **Adj. WF:** | n/a | - | 4.1e-4 | -0.001 | - | - | - | - | - | 2.799 | 0.015* |
| PD ALL | **BP:** | - | 0.018 | - | 0.072 | 0.025 | - | -0.079 | - | -0.022 | 16.14 | 6.02e-08*** |
| | **DL:** | - | - | 0.035 | -0.041 | - | - | -0.027 | - | - | 11.72 | 4.16e-6*** |
| | **WF:** | - | - | 0.041 | -0.054 | - | - | -0.018 | - | - | 10.16 | 6.33e-5*** |
| | **PC:** | - | - | -3.847 | 5.555 | - | -5.567 | 3.385 | - | - | 10.43 | 1.42e-5*** |
| | **BP Error:** | - | - | - | - | 0.002 | 0.009 | - | - | -0.002 | 7.031 | 8.39e-4*** |
| | **Adj. WF:** | - | - | - | - | 3.1e-4 | 6.4e-4 | - | - | -2.4e-4 | 3.261 | 0.033* |

*PD ON = Parkinson's disease (PD) patients on medication at time of time perception task. PD OFF = Parkinson's disease patients off medication at time of time perception task. PD ALL = All Parkinson's disease patients (both on and off medication state data) who completed the time perception task using medication (state) as an additional regressor. Dis Dur = PD Duration; Mult Meds = Multiple Dopaminergic Medications; LEDD = Levodopa Equivalency Daily Dose; Dep = Depression; UPDRS = United Parkinson's Disease Rating Scale; ICD = Impulse Control Disorder; QUIP = Questionnaire for Impulsive-Compulsive Disorders in Parkinson's Disease; Dx = Diagnosis. BP = Bisection Point; DL = Difference Limen; WF = Weber Fraction; PC = Percent Trials Correct. Significance is based on model significance, not individual variable significance. Significance based on p < 0.05*;0.01**;0.001****

Table S4: Doses and types of dopaminergic therapies (DT) of all PD subjects.

| ID | Carbidopa/Levodopa Dose (25/100mg) | Other DT | Levodopa Equivalent Dose (LED) | Levodopa Equivalent Daily Dose (LEDD) |
|---|---|---|---|---|
| 1 | 1 TID | 3mg Pramiprexole | 300mg QD | 600mg |
| 2 | 1.5 TID | 1mg Ropinirole | 20mg TID | 470mg |
| 3 | 1 6x/day | 100mg Amantadine | 100mg BID | 700mg |
| 4 | 2 TID | 200mg Entacapone | 133mg TID | 733mg |
| 5 | 1 TID | n/a | n/a | 300mg |
| 6 | 2 TID | n/a | n/a | 600mg |
| 7 | 1 QID | 0.25mg Ropinirole | 5 mg TID | 415mg |
| 8 | 1 TID | n/a | n/a | 300mg |
| 9 | 2 5x/day | n/a | n/a | 1000mg |
| 10 | 1 QID | 5mg Selegiline<br>4mg Ropinerole | 50mg QD<br>80mg BID | 610mg |
| 11 | 1 TID | 1mg Rasagiline | 100mg QD | 400mg |
| 12 | 2 1/2 TID | n/a | n/a | 750mg |
| 13 | 1.5 TID | n/a | n/a | 450mg |
| 14 | 1.5 7x/day | n/a | n/a | 950mg |
| 15 | 1 QD | 1.5mg Pramipexole | 150mg TID | 650mg |
| 16 | ½ TID | 100mg Amantadine | 100mg BID | 350mg |
| 17 | 1 TID | n/a | n/a | 300mg |
| 18 | 1 BID | 5mg Selegiline<br>100mg Amantadine | 5mg QD<br>100mg BID | 450 mg |
| 19 | 1 TID | 1mg Rasagiline<br>100mg Amantadine | 100mg QD<br>100mg BID | 600mg |

*For dosage frequency: QD = Once a day; BID = Twice a day; TID = Three times a day; QID = Four times a day. Subjects were asked to withhold all PD medications during off medication visits.*

Table S5: Beta coefficients from multivariate logistic regression models utilizing dimensions from principal component analysis (PCA) as independent predictors of dependent clinical features of PD presentation for both on and off medication groups.

| ***OFF MEDICATION*** | **ICD DX** | **DEP DX** | **DT** | **PD DUR** | **F-stat** | **p-value** |
|---|---|---|---|---|---|---|
| **Dimension 1** | -0.103 | -0.002 | -0.070 | 0.072 | 5.250 | 0.0184* |
| **Dimension 2** | 0.141 | 0.175 | -0.207 | -0.165 | 2.956 | 0.0815 |
| **Dimension 3** | -0.157 | -0.103 | -0.024 | -0.162 | 0.776 | 0.5673 |
| **Dimension 4** | 0.580 | -0.163 | 0.096 | 0.204 | 3.316 | 0.0625 |
| **Dimension 5** | 0.370 | 0.737 | 1.275 | 0.336 | 0.704 | 0.6084 |
| **Dimension 6** | -0.817 | -0.243 | 0.095 | -3.135 | 1.152 | 0.3927 |
| ***ON MEDICATION*** | **ICD DX** | **DEP DX** | **DT** | **PD DUR** | **F-stat** | **p-value** |
| **Dimension 1** | -0.032 | 0.062 | -0.030 | -0.004 | 0.617 | 0.6614 |
| **Dimension 2** | 0.218 | 0.184 | -0.124 | -0.160 | 6.874 | 0.0080** |
| **Dimension 3** | 0.157 | 0.052 | -0.073 | 0.234 | 0.670 | 0.6288 |
| **Dimension 4** | -0.582 | 0.394 | -0.542 | -0.971 | 1.953 | 0.1856 |
| **Dimension 5** | 0.430 | 0.363 | 0.340 | 0.139 | 0.045 | 0.9952 |
| **Dimension 6** | 0.611 | 0.603 | -0.334 | 1.293 | 0.197 | 0.9339 |

*ICD DX = Impulse Control Disorder Diagnosis; DEP = Depression; DT = Dopaminergic Therapies; PD DUR = Parkinson's disease duration. Significance based on* $p <$ *0.05*;0.01**;0.001**

Table S6: Results of Fischer Exact Tests to compare the categorical independent variable, Impulse Control Disorder (ICD), for Parkinson's Disease (PD) patients off medication based on the dependent variable, the Bisection Point (BP).

| Comparison Group | Odds Ratio | p-value |
|---|---|---|
| ICD Positive | 0.001 | 1.46e-5*** |
| ICD Negative | 1.279 | 0.7648 |
| Underestimate Intervals | 0.001 | 0.0115* |
| Overestimate Intervals | 0.322 | 0.0233* |

*Significance based on p < 0.05*;0.01**;0.001****

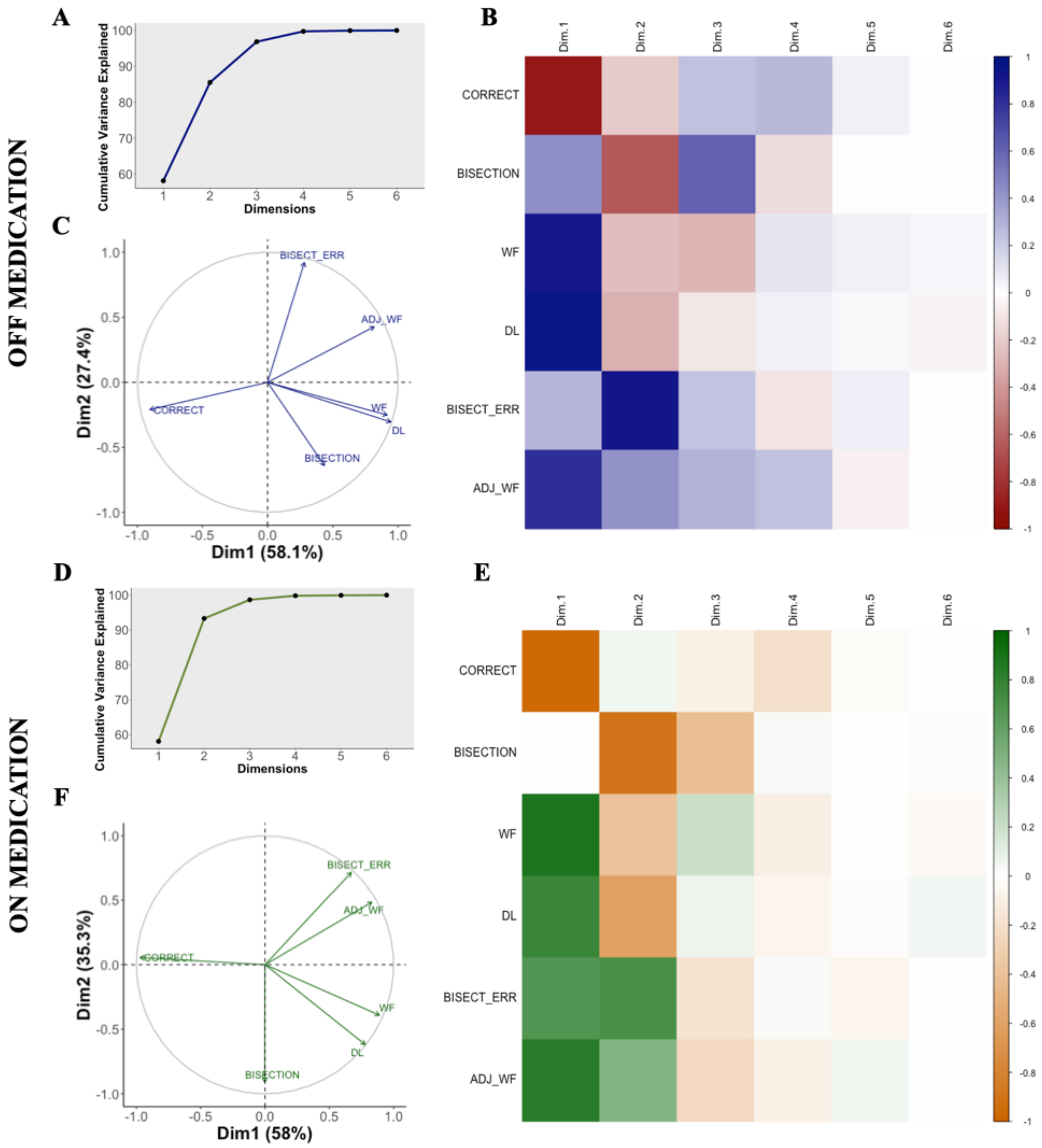


Figure S1: Dimension contributions from Principal Component Analysis (PCA) utilizing time perception measures for both on and off medication groups. A. Line plot showing the cumulative variance explained by each of six dimensions for the off medication Parkinson's disease group. B. Shows a correlation plot of the six dimensions and the six time perception measures for the off medication group, with a deeper blue indicating more positively correlated and a deeper red indicating more negatively correlated dimensions. White indicates little to no correlation. C. A variable plot for the off medication group. D. Line plot for the on medication Parkinson's disease group. E. Correlation heat map for the on medication group, with a deeper green indicating more positively correlated and a deeper orange indicating more negatively correlated. White indicates little to no correlation. F. Variable plot for the on medication group.

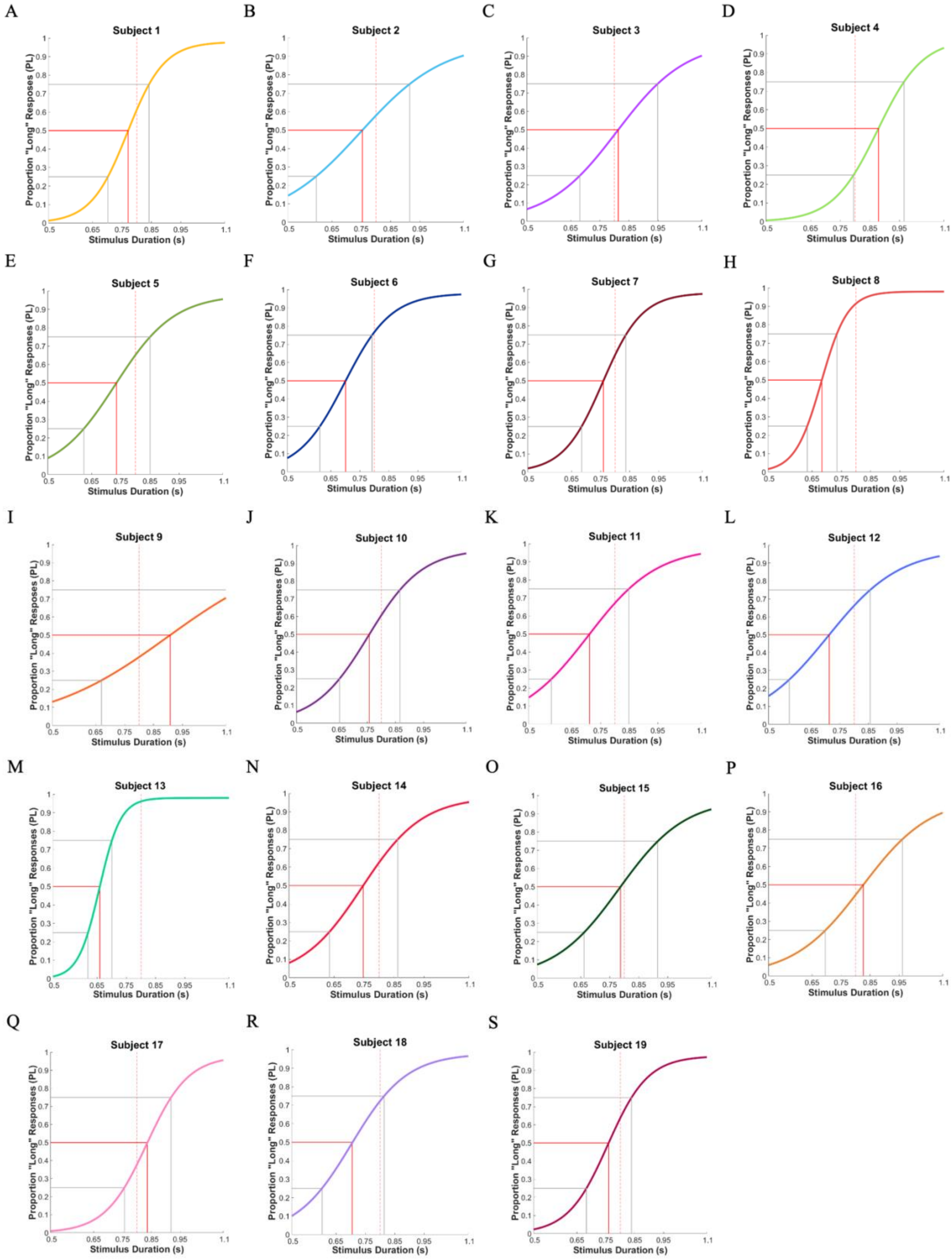
A
Subject 1
B
Subject 2
C
Subject 3
D
Subject 4
E
Subject 5
F
Subject 6
G
Subject 7
H
Subject 8
I
Subject 9
J
Subject 10
K
Subject 11
L
Subject 12
M
Subject 13
N
Subject 14
O
Subject 15
P
Subject 16
Q
Subject 17
R
Subject 18
S
Subject 19
Proportion "Long" Responses (PL)
Stimulus Duration (s)

Figure S2: Psychometric functions of interval timing from all 19 patients with Parkinson's disease. A-S. Individual psychometric function resulting from 500 trials (both sessions 1 and 2) of the temporal bisection task in the off medication state.
